# Decoding the underlying neural activity of neurodegeneration in Alzheimer’s Disease

**DOI:** 10.1101/2022.08.08.503255

**Authors:** Haolin Li, Hong Guo

## Abstract

Alzheimer’s Disease (AD) is the most prevalent chronic neurodegenerative disease, affecting more than 55 million people worldwide. AD is clinically characterized by a progressive loss of memory and other cognitive functions, which implicate the cholinergic system. Activity-induced neuroprotection based on gamma oscillation–audiovisual stimuli–is shown to be possible. Bulk RNA-seq data from treatment involving sensory stimuli were obtained, and bioinformatics analyses were performed. Activity-induced neuroprotection was found to be strongly associated with the downregulation of synaptic calcium signaling and cholinergic transmission. Candidate genes were screened in consultation with the Allen Brain Atlas Transcriptomics Explorer. We propose that Chrnb4 and Plcb4 are potential therapeutic targets for AD treatment. Chrnb4 has already been identified as a potential immune-relevant drug target gene using ontology inference, network analysis, and methylation signal, while multiple studies suggest that primary phospholipase C proteins play an integral role in inducing cell death in AD. Hence, both Chrnb4 and Plcb4 should be considered for future in vitro and in vivo studies as candidate genes.

## INTRODUCTION

Alzheimer’s Disease (AD) is the most prevalent chronic neurodegenerative disease, affecting more than 55 million people worldwide. It is the sixth leading cause of death in the United States and the fifth leading cause of death among Americans aged 65 and older [1]. AD is progressive and irreversible, clinically characterized by a progressive loss of memory and other cognitive functions–ultimately inducing dementia [2,3]. Although not fully understood, the pathophysiology of AD is largely represented by the neurotoxic events triggered by the beta-amyloid (Aβ) cascade and by cytoskeletal abnormalities subsequent to the hyperphosphorylation of microtubule-associated Tau protein in neurons. These processes lead to the formation of neuritic plaques [4] and neurofibrillary tangles [5], respectively, which are recognized as pathological hallmarks of AD [6].

Currently, pharmacological treatments for AD are largely limited to antidementia drugs–for example, memantine [7]. While clinical benefits have been observed, these treatments are limited to temporary, symptomatic support to cognitive functions while the disease progresses. Therapies targetting Aβ-plaques have been the focus of efforts to develop new pharmaceutical compounds with disease-modifying properties, but these approaches have failed to show any clinical promise [8]. On the other hand, neuroinflammation caused by microglial activation has emerged as an active area of research based on risk gene analysis [9,10], but there have yet to be any promising leads. Since AD is only 10% hereditary and 90% sporadic and since its exact cause remains unknown, there is no current method to prevent it.

The progressive cognitive decline of AD has been attributed to the observed cholinergic deficit [11]. Briefly, central cholinergic neurotransmission predominantly provides diffuse innervations, changes neuronal excitability, alters presynaptic release of neurotransmitters, and coordinates the firing of groups of neurons. In AD models, markers within cholinergic basal forebrain (CBF) neurons, which provide the major source of cholinergic innervation to the cerebral cortex and hippocampus, have been found to be downregulated; CBF cell loss and reduced cortical choline acetyltransferase activity were also noted [12,13,14]. Furthermore, CBF cortical projection neurons contain distinctive AD hallmarks and undergo neurochemical phenotypic alterations during disease progression [15]. In fact, delaying this neurodegeneration or minimizing its consequences is the mechanism of action for most currently available drug treatments for cognitive dysfunction in AD. Such cholinesterase inhibitors such as donepezil, rivastigmine, and galantamine reversibly inactivate cholinesterases, thereby inhibiting the hydrolysis of acetylcholine (ACh) and subsequently increasing ACh concentration in synapses. Moreover, since regions of the forebrain also contain non-cholinergic neurons–for instance, GABAergic interneurons–and since neuropeptides–including inhibitory neuropeptide galanin (GAL) Aβ–often colocalize with CBF neurons [16], it is therefore likely that understanding neurotransmitter interactions, in particular Ach, may provide novel targets for the development of potential treatments.

Alterations to neurochemical states result in changes to neural activity. Many studies have used techniques such as optogenetics and frequency entrainment to investigate the effect of certain sensory stimuli on neural activity and responses. To start, gamma oscillation has been implicated in cognitive processing [17,18,19]. Also, it is prominent in the hippocampus, suggesting its role in memory functions [20,21]. It is therefore not surprising that disorders characterized by memory impairment, such as AD, are associated with aberrant activity of gamma oscillations [22]. In fact, Klein et al. (2016) reported changes in gamma oscillations in the entorhinal cortex at an early stage of AD in animal model [23]. In 2016 and 2019, Tsai lab [45, 46, 47] demonstrated that 40Hz gamma oscillation entrainment through audiovisual stimuli ameliorates AD hallmarks and initiates activity-induced neuroprotection. They hypothesized that their results are due to alternative phenotypic activation of microglia. However, microglial activation must correlate with endogenous processes of neuronal cells. In this study, we analyzed RNA-seq data from Tsai lab [45, 46, 47] and screened for candidate genes using bioinformatics analyses. We aimed to uncover the mechanism underlying gamma oscillation entrainment treatment [45, 46, 47] and identify candidate genes with neuronal functions for further investigation. We hypothesized that cholinergic pathway genes aid in the modulation of neural activity, thereby granting activity-induced neuroprotection. By altering specific subunit interactions, we propose Chrnb4 and Plcb4 as potential therapeutic targets for AD treatment. Since audiovisual stimuli could be extended to incorporate music, we, here, present a way to quantify 40Hz content within any sound.

## METHODS

### Bioinformatics analyses

Bulk RNA-seq data were obtained from the Gene Expression Omnibus Accession Viewer (GSE77471). Bioinformatics analyses were performed using RStudio. All of the statistical details for each analysis were described in the figure legends. A volcano plot in the form of p-value versus magnitude of change was created using the stat package. Genes with large fold changes that are statistically significant were isolated. GO analysis was subsequently performed with the subset comprising the genes of interest using the gprofiler2 package, and enriched terms were visualized using ggplot2 and GOplot [24]. The genes in the two most significant GO terms were individually evaluated using the whole cortex and hippocampus single-cell RNA-seq data for mice from the Allen Brain Atlas Transcriptomics Explorer, from which mean neuronal expression levels were weighed against the types of cells with gene expression. 2 candidate genes were then selected.

### Soundwave quantification

R packages tuneR [25], signal [26], and oce [27] were used to cut wave files and build spectrograms. Using Fast Fourier Transform (FFT; an optimized algorithm for implementing Discrete Fourier Transformation), a sound sequence was sampled and divided into its spectral frequency components–single sinusoidal oscillations at distinct frequencies with unique amplitude and phase. The waveform was first plotted, and spectrogram parameters were then defined. After the code was functionalized, spectrograms were generated systematically.

## RESULTS

### Bioinformatics analyses show enrichment in synaptic nicotinic cholinergic transmission and cellcell signaling

To investigate the neuronal mechanism behind Tsai lab’s [45, 46, 47] treatment and to uncover potential drug treatment target genes or physical therapy causes, we analyzed their 40Hz gamma oscillation entrainment treatment group bulk RNA-seq data. We began by identifying statistically significant (adj p-value < 0.05) and differentially-expressed (log fold change > 1) genes using a volcano plot (Figure 1A). We found 198 such genes of interest. To interpret biological phenomena underlying 40Hz treatment, we then conducted Gene Ontology (GO) analysis and ranked the GO terms with respect to their negative log adj p-values. This way, we can interpret high throughput molecular data and define enriched gene sets. Here, we see enrichment in synaptic transmission and cell-cell signaling (Figure 1B). To remove redundant and extraneous terms (Figure 1C), we then congregated terms with overlaps greater than 75 percent.

**Figure 1.**
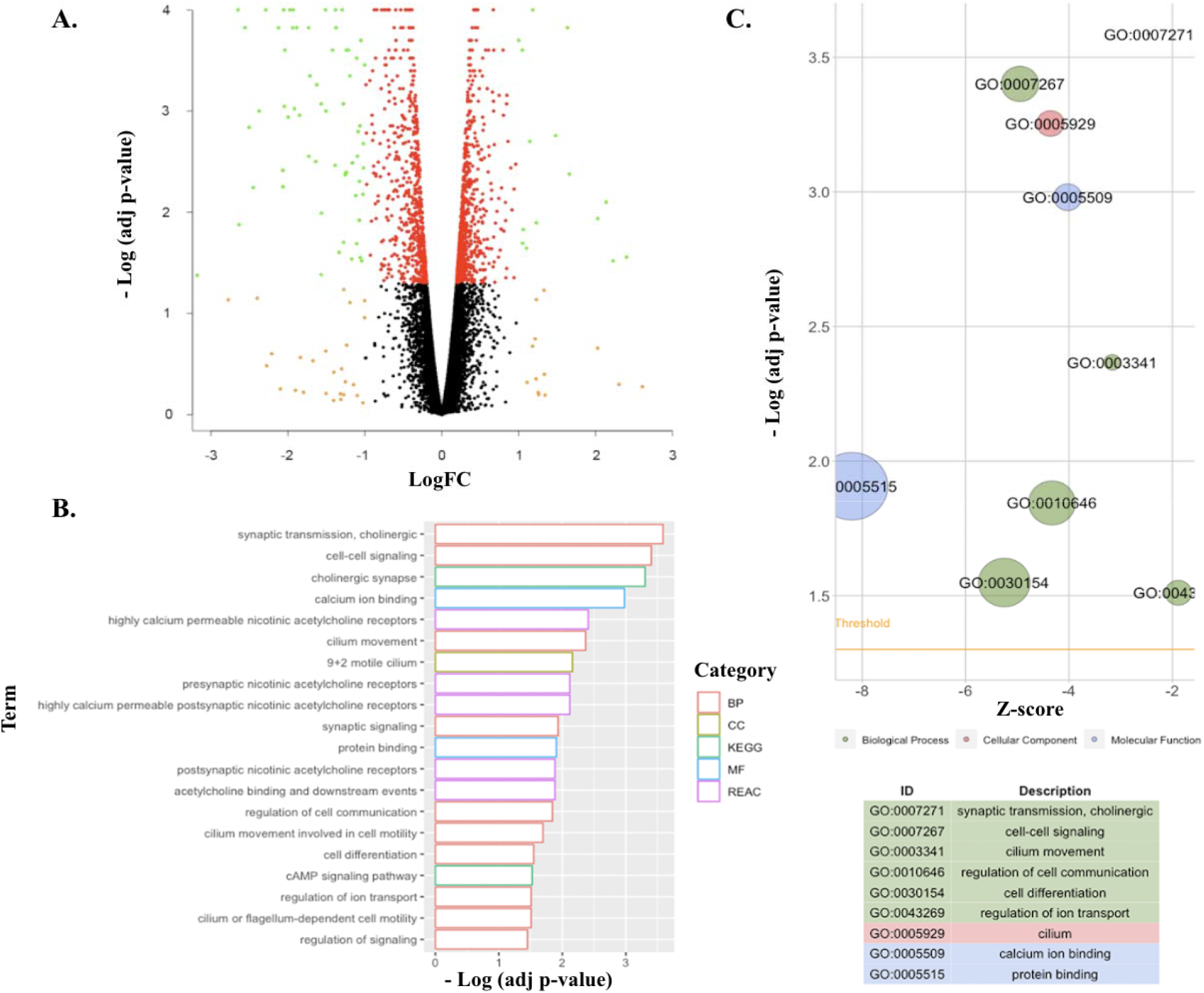
Analyses show enrichment in synaptic nicotinic cholinergic transmission and cell-cell signaling. A. Volcano plot determining the 198 genes of interest. Red: adj p < 0.05. Orange: |LogFC| > 1. Green: adj p < 0.05 and |LogFC| > 1. B. Bar plot ranking the GO terms by statistical significance (adj p < 0.05). C. Extraneous and redundant terms removed via term reduction and z-score added to predict regulation.

To better visualize significantly enriched terms, a circle plot (Figure 2A) was generated. Each dot, here, represents a specific gene, with blue representing downregulation and red representing upregulation. Most genes were downregulated in Tsai lab’s [45, 46, 47] treatment. The two most significant GO terms were synaptic nicotinic cholinergic transmission and cell-cell signaling, with adjusted p-values of 0.00026 and 0.00040 respectively.

**Figure 2.**
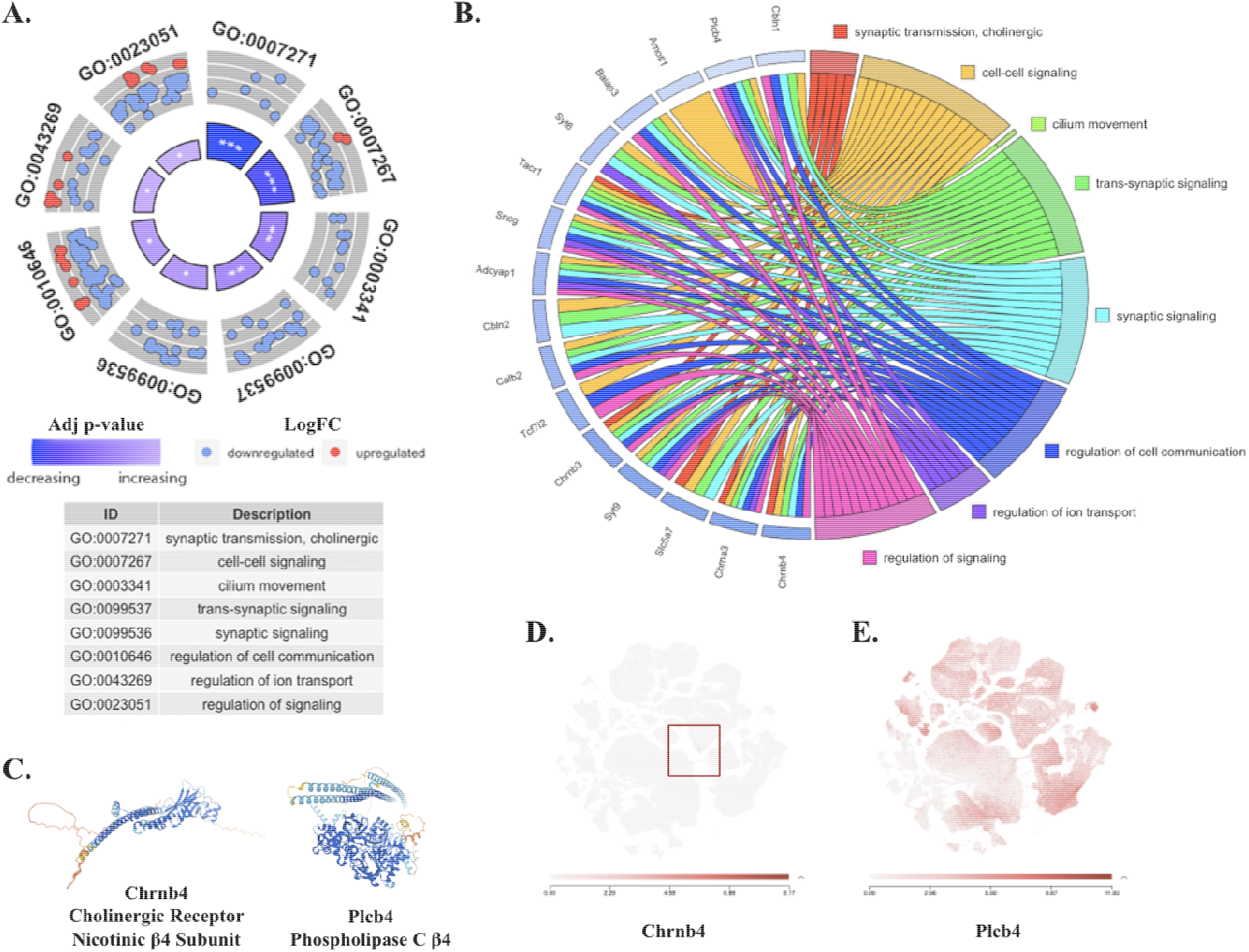
Identification of candidate genes from enriched GO terms. A. Visualization of individual differentially expressed genes in chosen GO terms. *** p < 0.001, ** p < 0.01, * p < 0.05. B. Selected candidate genes mapped. Blue outlines represent the degree of downregulation. Final candidate genes: Chrnb4 and Plcb4. C. Predicted protein structures of candidate genes. 2D-E. Neuronal expression levels of candidate genes.

### Identification of candidate genes from enriched GO terms

To identify potential therapeutic target genes for AD, we further analyzed genes constituting the two most significant GO terms. Specifically, the 6 genes belonging to synaptic cholinergic transmission and the 32 genes belonging to cell-cell signaling were individually evaluated in consultation with the Allen Brain Atlas. For quantification, we calculated a weighted score for each gene from their mean neuronal expression level and the number of cell subtypes expressing them. Because all 6 cholinergic-related genes were also present in the cell-cell signaling term, we ultimately chose 16 genes out of the total 32 genes for further analysis, and a chord plot (Figure 2B) was generated to depict their relationship with each other. In choosing our final 2 candidate genes, we evaluated these 16 genes by building a protein network of known and predicted interactions using STRING: functional protein association networks (Figure S1).

Chrnb4 (cholinergic receptor nicotinic β4 subunit) was chosen as our first candidate gene because it was the most downregulated after treatment (Figure 2B); Plcb4 (phospholipase C β4 isoform) was chosen as our second because of its sustained high neuronal expression levels in a wide array of cell types (Figure 2E). Their predicted protein structures (Figure 2C) and their neuronal expression level scatterplots (Figures 2D-E) were generated using AlphaFold and the Allen Brain Atlas, respectively. We note that, in normal mice, Chrnb4 is expressed in specific neuron subtypes, while Plcb4 is previously known to be ubiquitously expressed only in neurons.

### Soundwave quantification for future treatment design

Besides identifying potential therapeutic target genes, we could project Tsai lab’s [45, 46, 47] treatment with audiovisual stimuli onto music therapy, whereby pieces of music are chosen through quantification of their 40Hz content and utilized as a possible clinical intervention for AD. Therefore, an approach is needed to efficiently analyze and quantify 40Hz content within sound. To achieve this, spectrograms were generated to quantify the amplitude of sound waves over time. We chose to analyze a short 40 Hz test tone to verify the accuracy of our approach (Figure 3A). Then, a random piece of music was chosen to test the effectiveness of the approach in analyzing and distinguishing long, complex sound waves (Figure 3B). As Tsai lab [45, 46, 47] utilized 40Hz gamma oscillation entrainment, we, here, present a possible method to quantify the amount of 40Hz within any piece of music.

**Figure 3.**
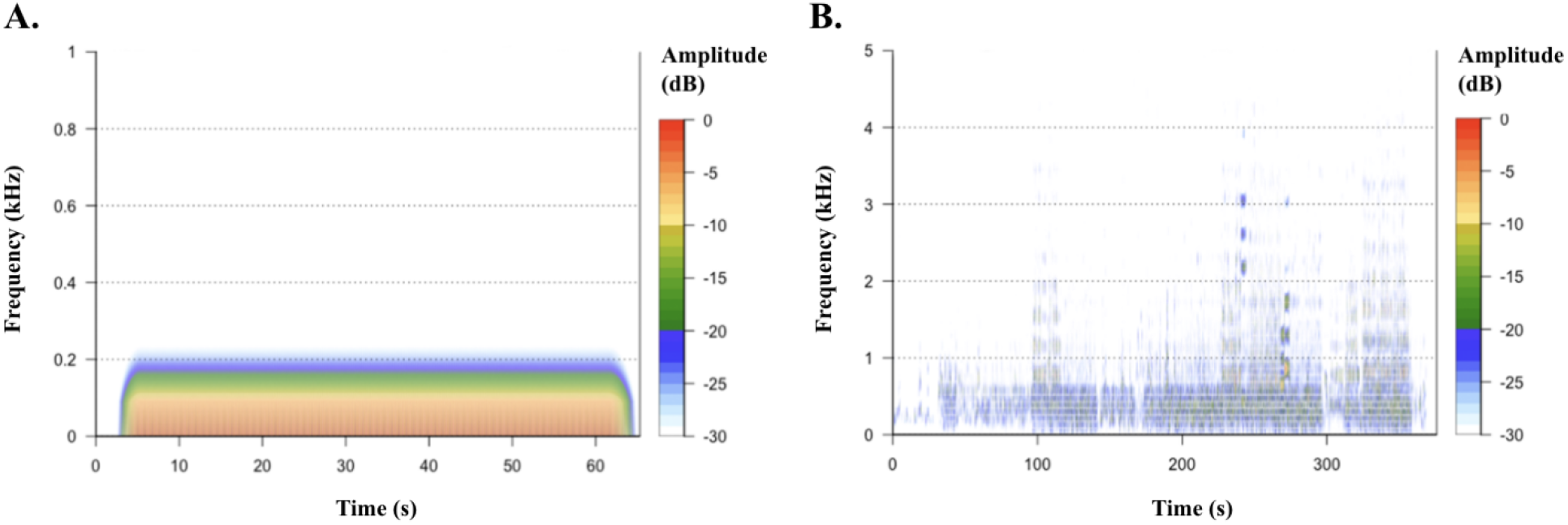
Soundwave quantification for future treatment design. A. 40 Hz test tone. B. Piece of music.

## DISCUSSION

In this study, we screened for candidate genes using comprehensive bioinformatics analyses. We concluded that activity-induced neuroprotection is strongly associated with the downregulation of synaptic calcium signaling and cholinergic transmission, specifically the nicotinic acetylcholine receptor (nAChR). We then isolated and identified Chrnb4 and Plcb4 as potential therapeutic targets for AD treatment. This study provided a glimpse into the underlying mechanisms of sensory stimuli and activity-induced neuroprotection. Our results may aid in the understanding of AD pathophysiology and contribute to the creation and translation of treatments, thereby helping individuals clinically and potentially improving patient outcomes.

Chrnb4, along with Chrna3 and Chrna5, constitute the important heteromeric 3β4 nAChR gene cluster. The nAChR subunits encoded by this locus form the predominant nicotinic receptor subtypes expressed in key central nervous system (CNS) sites, such as the medial habenula [28]. Plcb4 codes for the strictly neuronal expressing phospholipase C (PLC) β4 isoform, which mobilizes intracellular calcium by depletion of intracellular stores [29].

Because, as established, AD implicates the cholinergic system, it is therefore not surprising that our results indicate an association between AD treatment and restoration of cholinergic function. Chrnb4 of the 3β4 nAChR cluster has already been identified as a potential immune-relevant drug target gene using ontology inference, network analysis, and methylation signal [30,31]. Also, agonists of another nAChR, the homomeric □7 nAChR, have been shown to rescue Aβ-induced neurotoxicity and improve cognitive function in AD models [32]. Furthermore, this induced protection is blocked by an □7 receptor antagonist [33].

Regarding Plcb4, primary phospholipase C proteins PLCβ and PLCγ are activated primarily by neurotransmitters. Among primary PLC isozymes, PLCγ1, PLCβ1, and, more importantly, PLCβ4 are highly expressed and differentially distributed, suggesting a specific role for each PLC subtype in different regions of the brain. Primary PLCs are key enzymes in maintaining intracellular calcium concentration and sustaining signal, thereby controlling neuronal activity. Dysregulation of primary PLC signaling and disruption of calcium homeostasis have been linked to AD [34,35,36], with recent studies linking intracellular calcium overload to reduction in dendritic spines which, in turn, causes synaptic collapse and mediates neurodegeneration [37,38,39]. Yet, specific pathways and protein interactions remain unclear. For example, PLC-γ was abnormally accumulated in neurofibrillary tangles [40]. Moreover, PLCγ1 gene expression was enriched in AD patients [41], and PLC-γ protein level was significantly higher in the cytosolic fraction of AD cortical tissue than in control brains; however, its specific activity is compromised in AD brains [42], suggesting that its inactivation might be related to the pathophysiology of AD. Also, restoration of PLCβ1 activation ameliorated hippocampal longterm potentiation impaired by Aβ plaque accumulation [43], but further elevation by Aβ facilitates calcium overload (Park et al., 2022). Therefore, other genes encoding primary phospholipase C proteins–with emphasis on Plcb4–should be investigated as potential targets for protecting neurons against excitotoxicity in AD.

## ACKNOWLEDGEMENTS

This study would not have been possible without the guidance and support of Dr. Yi Lu. This research received no funding.

